# Compound interaction screen on a photoactivatable cellulose membrane (CISCM) identifies drug targets

**DOI:** 10.1101/2022.04.03.486868

**Authors:** Teresa Melder, Peter Lindemann, Alexander Welle, Vanessa Trouillet, Stefan Heißler, Marc Nazaré, Matthias Selbach

## Abstract

Identifying the protein targets of drugs is an important but tedious process. Existing proteomic approaches enable unbiased target identification but lack the throughput needed to screen larger compound libraries. Here, we present a compound interaction screen on a photoactivatable cellulose membrane (CISCM) that enables target identification of several drugs in parallel. To this end, we use diazirine-based undirected photoaffinity labeling (PAL) to immobilize compounds on cellulose membranes. Functionalized membranes are then incubated with protein extract and specific targets are identified via quantitative affinity purification and mass spectrometry. CISCM reliably identifies known targets of natural products in less than three hours of analysis time per compound. In summary, we show that combining undirected photoimmobilization of compounds on cellulose with quantitative interaction proteomics provides an efficient means to identify the targets of natural products.

---

Phenotypic screening has emerged as a key driver for biomedical innovation allowing the discovery of unknown therapeutic mechanisms of small molecules. A major challenge in such forward chemical genetic approaches is the chemoproteomics-based deconvolution and characterization of protein target and mode of action of the identified small molecule.^[1–3]^ Classical affinity-based target identification (ID) involves one distinct derivatization of the compound by one linker trajectory. This requires extensive efforts by structure activity relationship (SAR) studies to obtain suitable small molecular probes for affinity-based pulldown assays -- a tedious and time consuming-process which may even unintentionally exclude additional target proteins.^[4,5]^ This applies even more for large natural product derived molecules, where a synthetic access is not available or tractable. For kinase targets, an alternative method builds on affinity beads that are broadly specific for a wide range of cellular kinases. Using these beads in competition with free kinase inhibitors of interest in different concentrations enables the target ID.^[6,7]^ One drawback of this approach is its limitation to kinase inhibitors. More recent proteomic approaches like thermal proteome profiling (TPP) and limited proteolysis–small-molecule mapping (LiP-SMap) do not require compound tagging/immobilization.^[8,9]^ However, these methods require deep characterization of proteomic samples and therefore long mass spectrometric measurement times. Therefore, target ID studies based on TPP and LiP-SMap are typically limited to a single compound.

Undirected photocrosslinking is an attractive alternative to immobilize small molecules on an affinity matrix.^[10–12]^ The chemo- and site-nonselective nature of the photocrosslinking reaction leads to a distribution of differently tagged products for each small molecule with no prior derivatization required. This enables simultaneous and parallel immobilization of multiple small molecules in an array format. Such arrays can be probed with a single tagged protein, isolated or in a whole cell protein extract, to assess its interaction with multiple small molecules (multiple compounds, one candidate target protein).^[13]^ Photo-immobilized small molecules can also be used to fish for interaction partners in whole cell protein extracts followed by unbiased target ID.^[14–16]^ However, since distinguishing specific target proteins and non-specific contaminants is challenging, such target ID experiments were so far limited to single compounds (one compound, multiple target proteins). To the best of our knowledge, undirected photocrosslinking was not yet described for high-throughput target ID of multiple compounds in parallel (multiple compounds, multiple candidate target proteins).

Quantitative affinity purification combined with mass spectrometry (q-AP-MS) uses quantification to distinguish specific interaction partners and non-specific contaminants.^[17,18]^ This efficient and automated identification of specific interactors enables large-scale interaction screens. In this context, cellulose provides a cheap, easy to handle, lightweight and printable solid support platform for large scale interaction screens. For example, synthetic peptides immobilized on cellulose arrays via SPOT synthesis ^[19]^ can be used to screen for interacting proteins in whole cell extracts.^[20–23]^ On the one hand, the high local concentration of peptide ligands on the cellulose matrix and the mild washing conditions preserve even weak interactions, enabling interaction screens with high sensitivity. On the other hand, comparing protein abundance across different pull-downs can distinguish specific interactors from non-specific contaminants, which leads to high specificity.

We reasoned that photo-immobilizing small molecules on cellulose membranes should allow us to generate affinity matrices to probe the interaction of many proteins with multiple small molecules in parallel (multiple compounds, multiple target proteins). To this end, we devised a prototypic compound interaction screen on a cellulose membrane (CISCM) for complex natural products consisting of six steps easy to carry out (Figure 1). First, a cellulose membrane (CM) is functionalized with a photocrosslinker. Second, small molecules are spotted onto this membrane. Third, the spotted small molecules are immobilized via undirected photocrosslinking. Fourth, the membranes are incubated with protein extracts (e.g. whole cell lysates), followed by mild washing. Fifth, individual spots are excised, proteins are digested and analyzed by mass spectrometry using LC-MS gradients of 45 minutes per replicate. Finally, specific interaction partners of individual compounds are identified via label free quantification (LFQ).^[24]^

**Figure 1.**
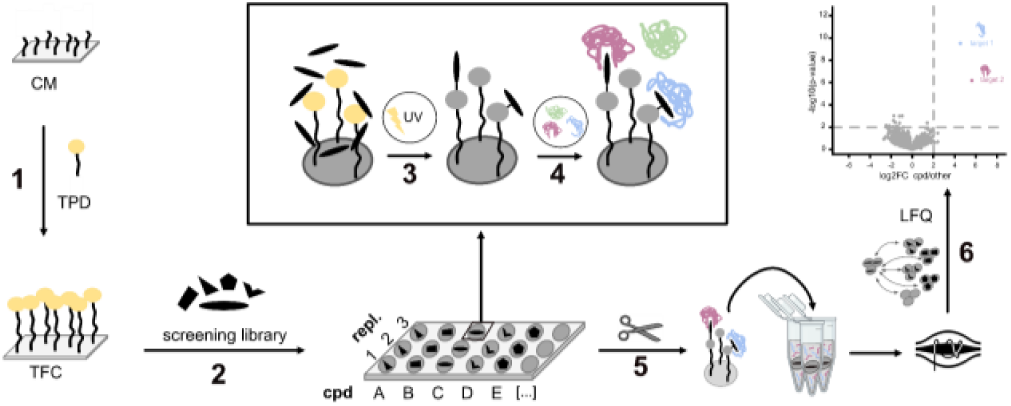
A compound interaction screen on a cellulose membrane (CISCM) in six steps. (1) Construction of a photoreactive cellulose membrane, (2) spotting of a screening library onto this membrane, (3) covalent attachment of physisorbed compounds via undirected UV-crosslinking, (4) pull-down of interacting proteins from a whole cell extract, (5) excision of each compound spot, protein digestion and LC-MS measurement, (6) target identification via quantitative analysis.

For undirected photocrosslinking we selected trifluoromethylphenyldiazirine (TPD) as a photo reactive precursor because of its superior crosslinking efficiency compared to other photocrosslinkers.^[25,26]^ We functionalized the cellulose membrane using a N-Hydroxysuccinimide (NHS)-based approach ^[27]^ and an amine-functionalized diazirine with a PEG-spacer (Scheme 1 a). This approach employs cellulose immobilized photogenerated carbene species that form covalent bonds with proximal small molecules by non-selectively inserting into carbon–heteroatom (C–Cl), heteroatom–hydrogen (O–H, N–H) and even carbon–hydrogen (Csp3–H, Csp2–H) bonds (Scheme 1b).

**Scheme 1.**
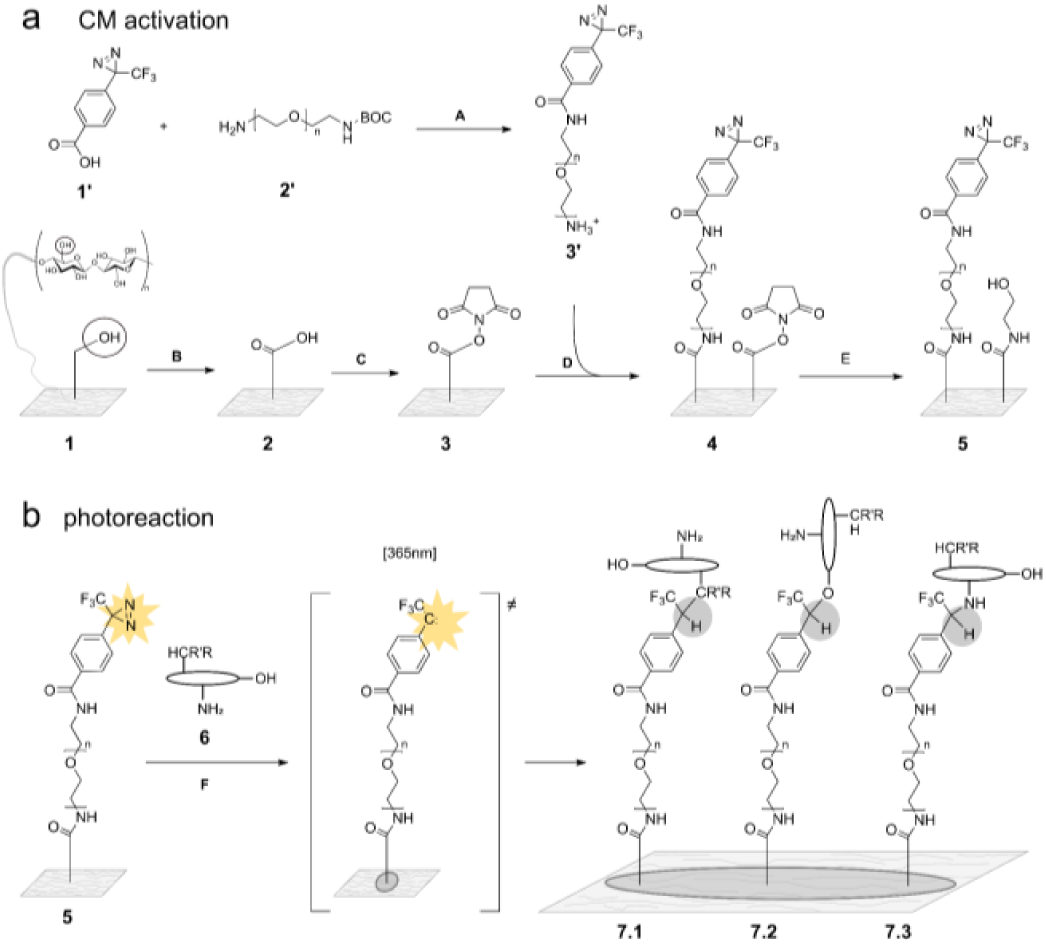
(a) Construction of a photocrosslinkable cellulose matrix: (top) Functionalization of 4-[3-(trifluoromethyl)-3H-diazirin-3-yl]benzoic acid (TDBA) (**1’**) with an amine-reactive PEG-spacer of variable length using Boc-NH-PEGn-CH2CH2NH2 (**2’**) to create a primary amine-containing trifluoromethylphenyldiazirine (TPD) (**3’**). (bottom) Stepwise activation of cellulose membranes (**1**) by TEMPO-mediated oxidation to get oxidized cellulose (**2**) and activation with NHS and 1-Ethyl-3-(3-dimethylaminopropyl)carbodiimide (EDC) to obtain NHS-activated cellulose (NAC) (**3**). Immobilization of the photocrosslinker (**3’**) on the activated cellulose (**3**) to form the TPD-functionalized cellulose (TFC) (**4**). Blocking of unreacted NHS-groups to obtain a blocked TFC membrane (**5**). (b) reaction principle of undirected photocrosslinking: spotting of constructed affinity matrix (**5**) with a compound solution of interest, here represented by a fictional compound (**6**), evaporation of the solvent, formation of reactive carbene induced by irradiation with UV-light at 365 nm, formation of different reaction products (**7.1, 7.2, 7.3**) of undirected photocrosslinking reaction between photoactivated 5 and dried compound of interest (**6**). Reaction conditions: (**A**) 1 eq. TDBA, 1.25 eq. Boc-N-amido-PEG-amine, 0.35 eq. 4-Dimethylaminopyridine (DMAP), 30 min, THF; 1.75 eq. EDC, 18 h, RT, dark; (**B**) NaOH (w(NaOH) = 10%), 18 h, H_2_O, RT; 0.39 mmol TEMPO, 14.1 mmol NaBr, 17.0 mmol NaOCl, 1 h, H_2_O, pH 10; (**C**) 30 mmol NHS, 0.4 mol/l EDC, sodium acetate, 1.5 h, H_2_O, pH 5; (**D**) 10 mM amine-PEG TPD, 21 h, THF, RT; (**E**) 1-3 M ethanolamine (EA), 1-2 h, H_2_O; (**F**) 10 mM compound solution, drying, irradiation at 365 nm.

To confirm the efficient functionalization of the cellulose we followed each of the functionalization steps shown in Scheme 1a (bottom) using X-ray photoelectron spectroscopy (XPS, Figure 2a) and attenuated total reflection Fourier-transformed infrared (ATR-FTIR) spectroscopy (Figure 2b, Supporting Figure 1). Comparison of high resolution C 1s spectra of unmodified (**1**) versus oxidized cellulose (**2**) shows the appearance of a peak of weak intensity at 289.2 eV binding energy, which corresponds to the newly formed carboxylic group in **2**. The formation of a carboxylic group could also be observed by an ATR-FTIR signal for carboxyl vibrations at 1599 cm^-1^ for the oxidized cellulose (**2**). Subsequent functionalization of the oxidized cellulose with EDC and NHS led to the detection of N 1s at 402.0 eV binding energy corresponding to the succinimide nitrogen bound to the oxygen of the ester group (1.0 atomic percent).^[28]^ A further species at 400.0 eV can be explained by the remaining of EDC which contains N=C=N groups and amine (5.6 atomic percent). The functionalization of cellulose with NHS could also be detected in ATR-FTIR showing vibrations for the amide group (1705 cm^-1^). The functionalization of NHS-activated cellulose (**3**) with a PEG-4 diazirine photocrosslinker led to a significant decrease of this ATR-FTIR signal and showed a signal at 688.2 eV binding energy in the high-resolution F 1s photoelectron spectrum corresponding to covalently bound fluorine atoms of the trifluoromethyl group. Both XPS and ATR-FTIR data therefore confirm a successful attachment of **3’**. The decreased signal of amide vibrations at 1705 cm^-1^ completely disappeared after blocking the membrane with ethanolamine (EA). As expected, the additional functional group introduced by EA furthermore led to a broad signal from 1670-1540 cm^-1^, instead of two separated peaks at 1650 cm^-1^ and 1539 cm^-1^.

**Figure 2.**
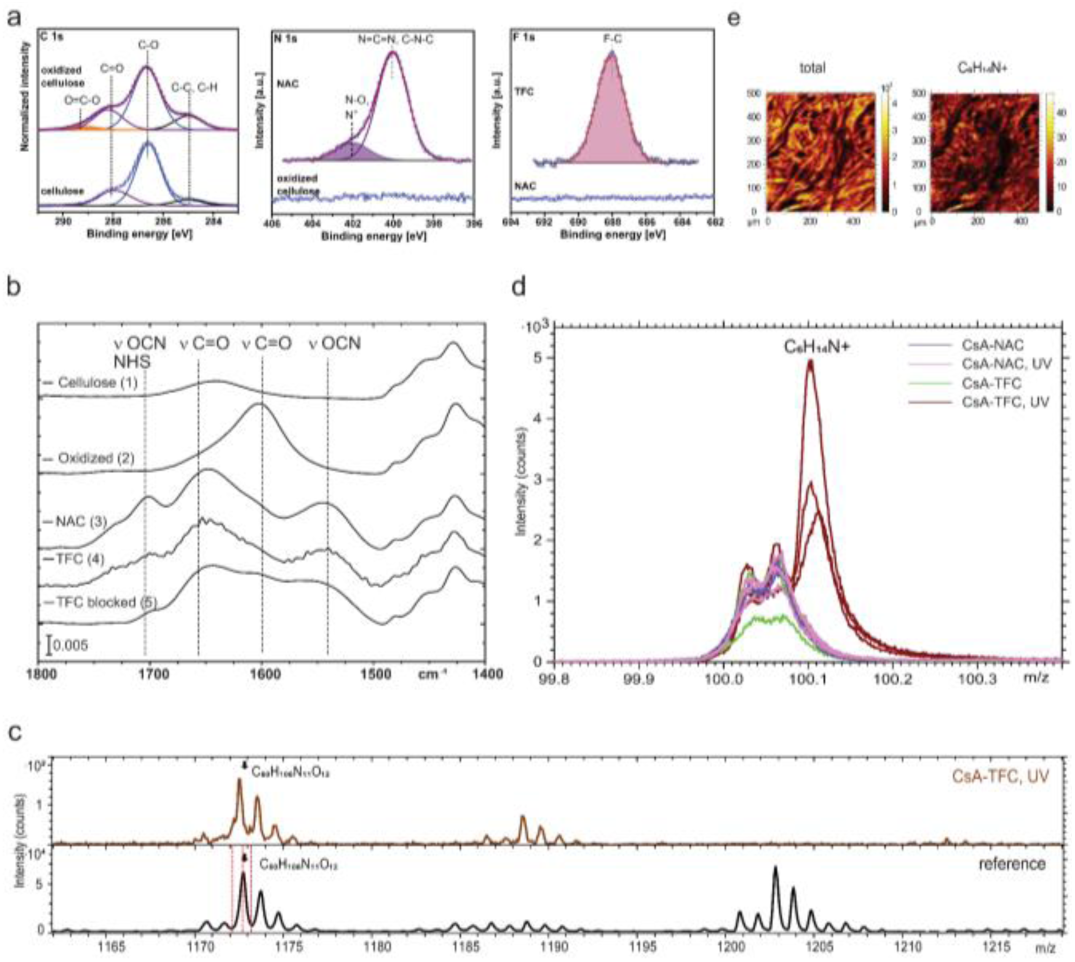
XPS, ATR-FTIR and ToF-SIMS spectra of stepwise cellulose functionalization. (a) C 1s XPS spectra for unmodified and oxidized cellulose (left), N 1s photoelectrons for oxidized and NHS-activated cellulose (NAC, center) and F 1s photoelectrons of NHS-activated and TPD-functionalized cellulose (TFC, right). (b) Selected region of ATR-FTIR spectra of unmodified and functionalized cellulose. (c) ToF-SIMS spectra of a CsA reference sample (drop-cast, bottom) and CsA-spotted TFC after UV irradiation (top) over a mass range of m/z 1160-1220. (d) ToF-SIMS spectrum in the mass range of the amino acid fingerprinting signal of N-methylated Leu (C_6_H_14_N^+^, m/z 100) across different samples: CsA spotted NAC (NHS-activated cellulose, purple), CsA-spotted UV-treated NAC (pink), CsA spotted TFC (green) and CsA spotted TFC and treated with UV (brown). Spectra were acquired at three different lateral positions across the corresponding sample. (e) ToF-SIMS chemical mapping showing the lateral distribution of different CsA-fragment signals across CsA spotted TFC irradiated with UV.

To evaluate if the TFC matrix allows photocrosslinking of small molecule drugs we first performed experiments using cyclosporine A (CsA) as a large polyfunctional natural product, (Figure 2c-e) using Time-of-Flight Secondary Ion Mass Spectrometry (ToF-SIMS) since it allows detection of both intact molecules and corresponding fragments. CsA was spotted in duplicates on a TPD functionalized cellulose (TFC) membrane and a NHS-activated cellulose (NAC) membrane as a control. Immobilization of CsA was induced by photocrosslinking for one replicate of each sample whereas the second replicate was kept in the dark. After intense washing, the resulting four samples were analyzed by ToF-SIMS. A drop cast sample of CsA was measured as a reference. The molecular ion of CsA indicating non-covalent attachment was only detected in the reference but could not be observed in any of the 4 samples (Figure 2c). A corresponding fragment ion of CsA (C_60_H_106_N_11_O_12_ ^+^) observed in the reference sample could only be identified in the CsA-spotted and UV-treated TFC sample, but not in any of the 3 control samples (Supporting Figure 2). Amino acid fingerprinting revealed a N-methylated leucine ion (C_6_H_14_N^+^) that could only be identified in CsA-spotted and UV-treated TFC but not in any of the 3 control samples (Figure 2d). These findings show that the covalent attachment of CsA to the cellulose membrane by a photoreaction requires both a diazirine photocrosslinker as well as the activation by UV-light. The covalent photoimmobilization of CsA was furthermore confirmed to be laterally homogeneous across the cellulose membrane and also detectable on the side of the membrane facing away from the UV light source during irradiations shown by chemical mapping with ToF-SIMS (Figure 2e).

Having shown that our approach can photoimmobilize small molecules onto a cellulose matrix, we next tested if this platform provides a suitable affinity matrix to assess protein-small molecule interactions. To this end, we selected cyclosporin A (CsA), tacrolimus (FK506) and sirolimus (rapamycin) as model compounds because of their well characterized protein binding partners and their high structural complexity.^[8,29–31]^ We also included lenalidomide as a member of the group of immunomodulatory drugs (IMiDs) that can pull-down their target ligand cereblon.^[32]^ All compounds were spotted onto the TFC membrane (**4**) in triplicates and immobilized via photocrosslinking. Unmodified, oxidized and TPD-activated cellulose without spotted compounds were used as controls. To obtain protein extracts we lysed Jurkat cells in lysis buffer. For the interaction screen, cellulose membranes were incubated with the lysate for two hours at 4 °C. After three washes with a detergent-free lysis buffer, membranes were air-dried. Cellulose spots corresponding to individual photocrosslinked compounds were excised and transferred into 96-well plates containing digestion buffer and processed for shotgun proteomic analyses using standard methods. All 21 samples (four compounds and three controls, in triplicates) were analyzed by high resolution LC-MS/MS on a Q Exactive HFX mass spectrometer.

Data analysis with MaxQuant ^[33]^ identified 3,383 protein groups (protein and peptide FDR of 1%) in all samples combined (Supplemental Table S1). To identify the proteins interacting specifically with a given immobilized compound we used label-free quantification (LFQ).^[24]^ Hierarchical clustering of differentially abundant proteins (ANOVA, FDR 5%) revealed clustering of replicate samples (Figure 3 a). To identify specific targets we compared protein abundances in the three replicates for a given compound to all other samples using the Student’s t-test and presented the data as volcano plots (Figure 3 b-e). As expected, the vast majority of identified proteins did not show preferential binding and can thus be considered non-specific background proteins. We selected specific binders requiring t-test p-values < 0.01 and fold changes of at least 4. For cyclosporine, this identified PPIF, PPIA, PPIL1 and PPIB as key specific interactors, corroborating previous results.^[29,34,35]^ Similarly, for immunosuppressant drugs Tacrolimus (FK506) and Sirolimus (Rapamycin) members of the FK506-binding protein (FKBP) family could be identified as specific binders. Importantly, all of the proteins identified as specific binders of these three natural products are previously known targets. Hence, CISCM can detect targets of natural products with high specificity. In contrast, we failed to detect relevant targets for the small drug lenalidomide, even though targeted immobilization of IMiDs can enrich their target cereblon ^[32]^, which is expressed in Jurkat cells we used.^[36]^ Our data therefore indicate that undirected photocrosslinking may not be efficient enough for fragment-like small molecules like lenalidomide only exhibiting a limited number of reactive attachment sites.

**Figure 3.**
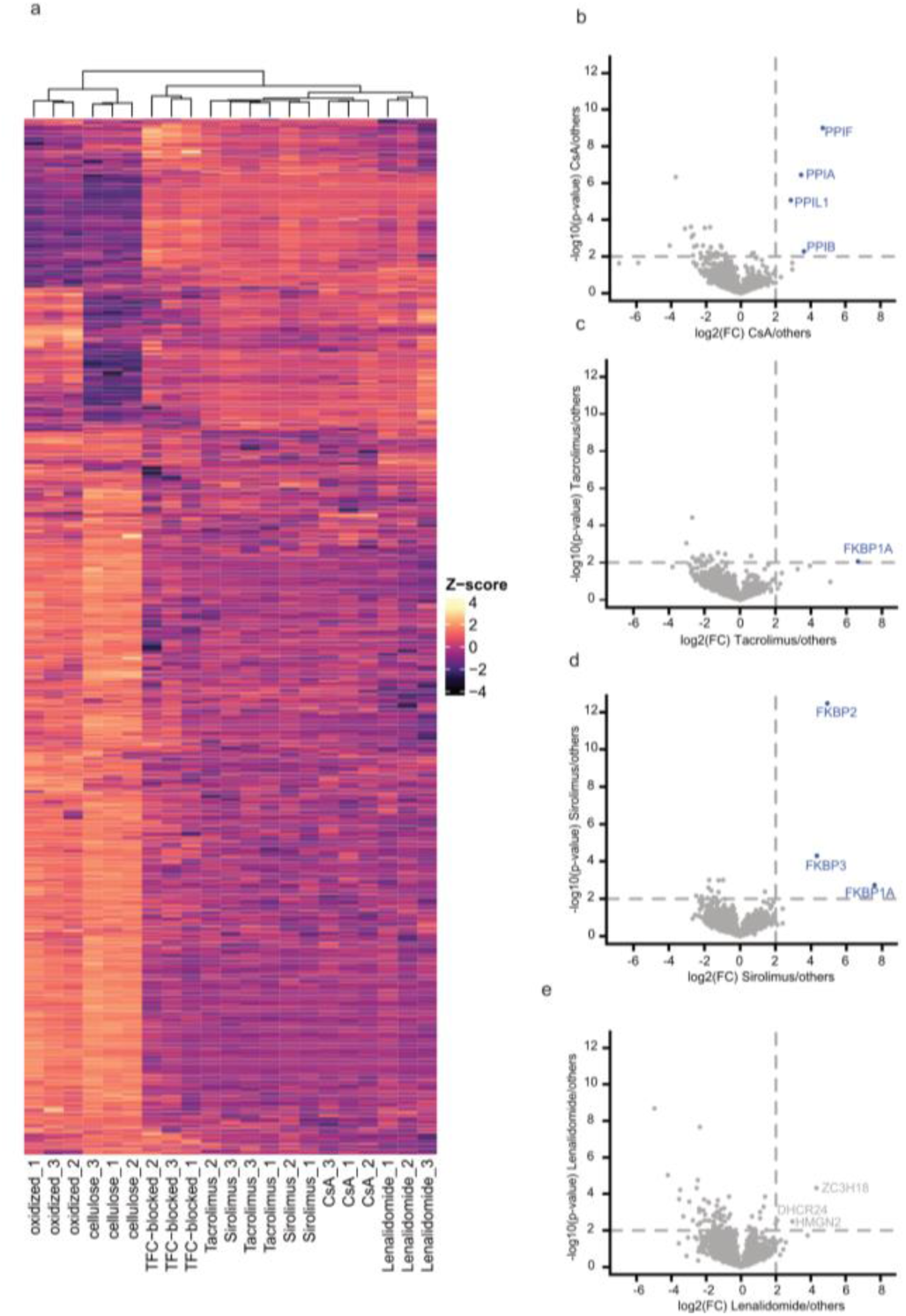
Differential protein identifications derived from AP-MS data. (a) Heatmap of Z-scores computed for ANOVA significant (FDR 5%, 250 randomizations) protein identifications of 4 immobilized compounds and 3 controls. Each column is an individual replicate of the total 21 samples. (b-e) Volcano plots displaying the log2 fold change (x-axis) against the Student’s t-test derived -log10 p-value (y-axis) for pairwise comparison of grouped triplicates of (b) cyclosporine, (c) Tacrolimus, (d) Sirolimus and (e) Lenalidomide, respectively, against all other samples. Proteins with t-test p-values < 0.01 and fold changes of at least 4 are labeled and known protein targets marked in blue.

In summary, we present a compound interaction screen on a photoactivatable cellulose membrane (CISCM) that is able to rapidly screen for drug targets in a parallel fashion. A key advantage of the approach is that it does not require tedious functionalization and previous SAR studies for linker implementation. While undirected photocrosslinking has been used before to assess interaction of individual candidate proteins with drug libraries, we show that q-AP-MS enables unbiased proteome-wide screening. Compared to immobilization-free methods like TPP and LiP-SMap that require long LC-MS measurement times, the high throughput of CISCM enables multiplexed analysis of several compounds in parallel. Current limitations of our method are due to the known limitations of diazirine-based photocrosslinking strategies. For example, the approach does not seem to work efficiently for small fragment-like compounds. However, its ability to reliably detect targets of larger and complex natural products makes it particularly suitable for this important class of drug compounds. Given its simplicity and throughput, CISCM is an attractive method, complementing existing approaches in particular in the context of drug leads derived from natural products.^[37,38]^

## Acknowledgements

The authors would like to thank Martha Hergeselle and Christian Sommer (MDC) for help with cell culture and mass spectrometry, respectively. We also thank Peter Schmieder (FMP) and members of the MDC proteomics community (Philipp Mertins lab, Ilaria Piazza lab, Fabian Coscia lab, Matthias Selbach lab) for fruitful discussions.

## Entry for the Table of Contents

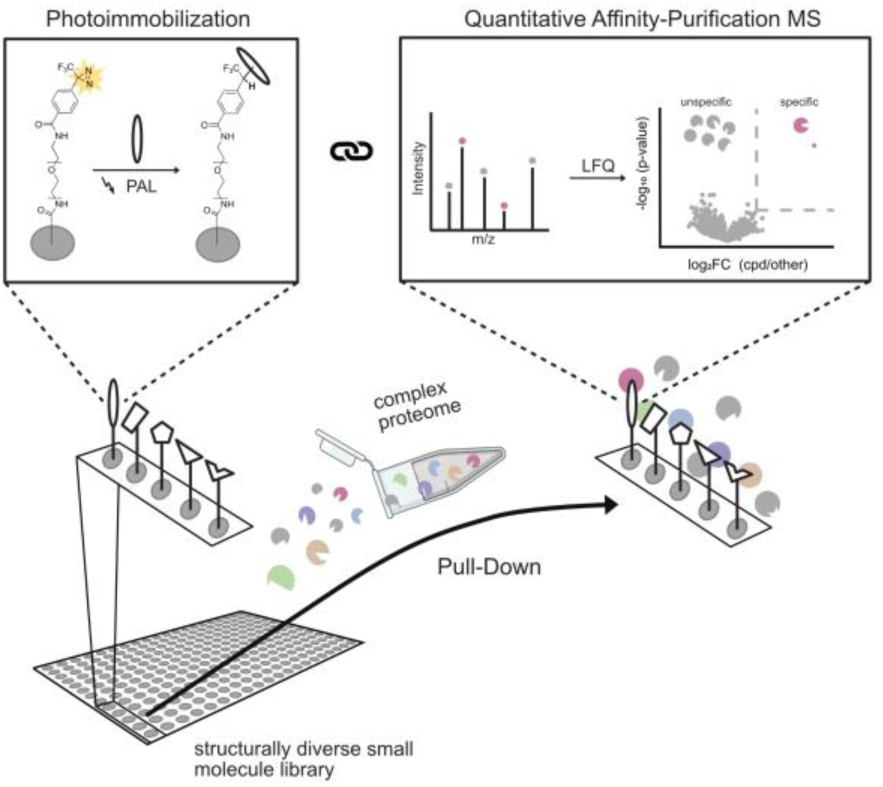

We present a new compound interaction screen on a cellulose membrane (CISCM) that combines undirected photoimmobilization of compounds on cellulose with affinity purification mass spectrometry and label free quantification (LFQ) to identify the protein targets of several compounds in parallel.

Institute and/or researcher Twitter usernames: @SelbachLab

