## Supporting Information for "Compound interaction screen on a photoactivatable cellulose membrane (CISCM) identifies drug targets"

##### Materials and Methods

1. General Methods

Commercial chemicals from Sigma Aldrich, Chess, Merck, TCI, Acros and Roche were used as supplied. In all reactions described in section 2-3 deionized water was used. For HPLC of synthesis products milli-Q water was used. For affinity purification-mass spectrometry (AP-MS) sample preparation LC-MS grade solvents were used. For modification filter membranes from Whatman type 50, 55 mm were used. Nuclear Magnetic Resonance (NMR) mono-(1H, 13C and 19F) were recorded on a Bruker AVANCE 300 and Bruker AV 600 MHz spectrometers. All 13C-NMR-spectra were recorded with 1H-broad-band decoupling. All chemical shifts are reported in ppm relative to tetramethylsilane ($\delta$ = 0.00 ppm) and were calibrated with respect to their respective deuterated solvents. Mass spectrometry analyses on synthesized structures (section 2) were performed with two different spectrometers using the same column and method: Column: Thermo Accuore RP-MS; Particle Size: 2.6 µm; Dimension: 30 × 2.1 mm; Eluent A: Water with 0.1% TFA; Eluent B: Acetonitrile with 0.1% TFA; Gradient: 0.00 min 95% A, 0.2 min 95% A, 1.1 min 1% A, 2.5 min Stoptime, 1.3 min Posttime; Flow rate: 0.8 ml/min; UV-detection: 220 nm, 254 nm, 300 nm. High resolution mass spectrometry analyses were carried out using the spectrometer Agilent Technologies 6220 Accurate Mass ToF LC/MS linked to Agilent Technologies HPLC 1200 Series, whereas liquid chromatography-mass spectrometry analysis was carried out employing Agilent Technologies 6120 Quadrupole LC/MS linked to Agilent Technologies HPLC 1290 Infinity. Thin Layer Chromatography was carried out on TLC-plates from Merck (Silicagel 60, fluorescence-indicator F254, layer thickness = 0.25 mm). For flash chromatography purifications, a Biotage Isolera One apparatus with RediSep®Rf Columns from Teledyne Isco, was used. Preparative HPLC: HPLC analyses were performed on a Shimadzu HPLC system, consisting of a CBM-20A controller, LC-20AP pump, SPD-20 a UV detector and a FRC-10 fraction collector. The separations were performed on a Macherey-Nagel VP250/21 Nucleodor 100-7 C18ec. Compounds were eluted at a flow rate of 30 ml/min, using water (Solvent A) and acetonitrile (Solvent B) as a mixture of solvents. AP-MS samples were analyzed by high resolution LC-MS/MS on a Thermo Fisher Q Exactive HFX mass spectrometer connected to a Thermo Fisher EASY-nLC 1200 System using a short gradient (45 min) on a reverse phase column (C18, particle size: 1.9 µm) and data dependent acquisition (DDA, Top20, MS2-resolution: 15K).

#### 2. Synthesis of a photoactive linker

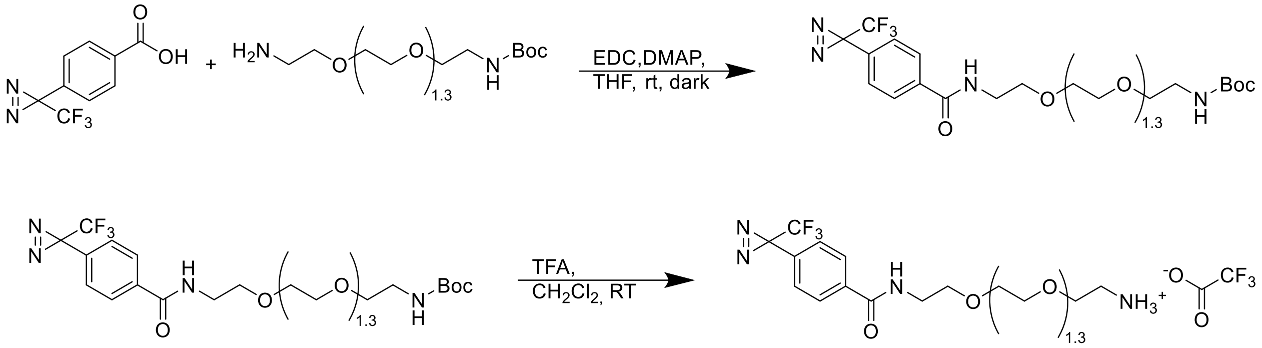


General procedure I: To a stirred solution of 4-[3-(trifluoromethyl)-3H-diazirin-3-yl]benzoic acid (TDBA) (1.00 eq.), Boc-N-amido-PEG-amine (1.25 eq.), and 4-Dimethylaminopyridine (DMAP) (1.217 mmol; 0.35 eq.) THF was added after 30 min. 1-(3-Dimethylaminopropyl)-3-ethylcarbodiimide (EDC) (1.75 eq.) was added and the reaction mixture was stirred for additional 18 hours under protection against light. After successful reaction the solvent was evaporated under reduced pressure. The crude product was purified by silica gel column flash chromatography using dichloromethane and methanol as solvents.

General procedure II: The boc protected amine was deprotected by treating the corresponding Boc-N-amido-PEG-TDBA solved in dichloromethane with trifluoro acidic acid at room temperature for 90 minutes under protection from light. After complete reaction the mixture was quenched with ice water and extracted two times with dichloromethane. The combined organic phases have been dried over magnesia sulfate and the solvent was evaporated under reduced pressure. The crude product was purified with preparative HPLC.

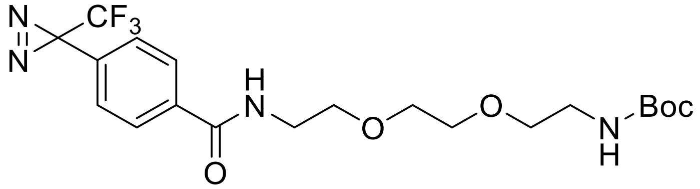

PEG-2 linker:

**tert-butyl (2-(2-(2-(4-(3-(trifluoromethyl)-3H-diazirin-3-yl)benzamido)ethoxy)ethoxy) ethyl)carbamate:[1]** To a solution of 100 mg TDBA (0.434 mmol; 1.00 eq.), and 423 µl N-Boc-2,2′-(ethylenedioxy)diethylamine (1.78 mmol; 4.1 eq.) as corresponding Boc-PEG-amine, and 18.6 mg DMAP (0.152 mmol; 0.35 eq.) in 6 ml THF, 146 mg EDC (0.760 mmol; 1.75 eq.) was added after 10 min. Following the general procedure I tert-butyl (2-(2-(2-(4-(3-(trifluoromethyl)-3H-diazirin-3-yl)benzamido)ethoxy)ethoxy)ethyl)carbamate (138 mg, 0.43 mmol, 69%) was obtained as an colorless oil.

1H NMR (600 MHz, DMSO-d6) δ in ppm 8.66 (t, J = 5.6 Hz, 1H), 7.98 – 7.92 (m, 2H), 7.37 (d, J = 8.1 Hz, 2H), 6.75 – 6.70 (m, 1H), 3.55 – 3.47 (m, 6H), 3.41 (q, J = 5.8 Hz, 2H), 3.36 (t, J = 6.2 Hz, 2H), 3.04 (q, J = 6.0 Hz, 2H), 1.36 (s, 9H). 13C NMR (151 MHz, DMSO) δ 165.2, 155.5, 149.3, 136.0, 130.1, 128.1, 126.4, 106.7, 77.5, 69.5, 69.4, 69.1, 68.7, 28.2. Calcd .mass for $C_{20}H_{28}F_{3}N_{4}O_{5}$, m/z 461.2006 $(M+H)^{+}$, found 461.2008.

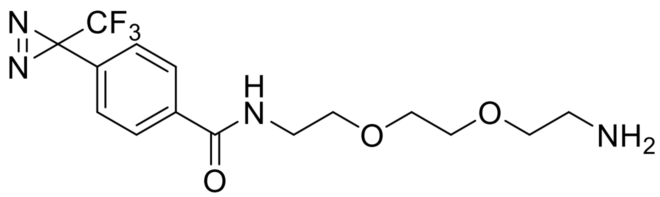


**N-(2-(2-(2-aminoethoxy)ethoxy)ethyl)-4-(3-(trifluoromethyl)-3H-diazirin-3-yl)benzamide:**[1] Starting with 137 mg procedure tert-butyl (2-(2-(2-(4-(3-(trifluoromethyl)-3H-diazirin-3-yl)benzamido)ethoxy)ethoxy)ethyl)carbamate (0.298 mmol; 1.00 eq.) in 5 ml dichloromethane and using 745 µl TFA (9.67 mmol; 32.5 eq.), following general procedure II N-(2-(2-(2-aminoethoxy)ethoxy)ethyl)-4-(3-(trifluoromethyl)-3H-diazirin-3-yl)benzamide (80.6 mg, 0.224 mmol, 81%) was obtained as TFA salt and colorless oil.
1H NMR (600 MHz, DMSO-d6) δ 8.68 (t, J = 5.7 Hz, 1H), 7.97 – 7.92 (m, 2H), 7.78 (s, 3H), 7.38 (d, J = 8.1 Hz, 2H), 3.57 (d, J = 4.5 Hz, 6H), 3.54 (t, J = 6.1 Hz, 2H), 3.43 (q, J = 6.0 Hz, 2H), 2.95 (q, J = 5.1 Hz, 2H). 13C NMR (151 MHz, DMSO) δ 165.2, 135.9, 130.2, 128.1, 126.4, 121.74 (q, J = 274.6 Hz), 69.7, 69.4, 68.7, 66.6, 40.1, 38.6, 28.03 (q, J = 39.8 Hz).
Calcd. mass for $C_{15}H_{20}F_3N_4O_3$, m/z 361.1482 $(M+H)^+$, found 361.1484.

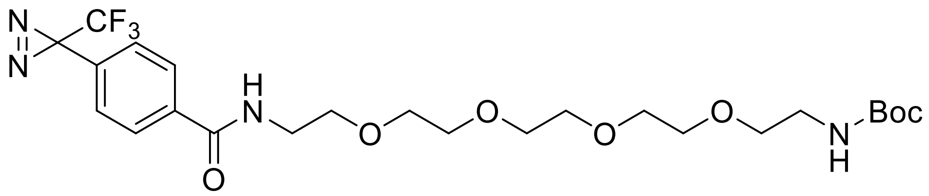


PEG-4 linker: tert-butyl **(1-oxo-1-(4-(3-(trifluoromethyl)-3H-diazirin-3-yl)phenyl)-5,8,11,14-tetraoxa-2-azahexadecan-16-yl)carbamate:** To a solution of 800 mg TDBA (3.48 mmol; 1.00 eq.), and 1.38 ml t-Boc-N-amido-PEG4-amine (4,34 mmol; 2,50 eq.) as corresponding Boc-PEG-amine, 148 mg DMAP (1.217 mmol; 0,35 eq.) in 50 ml THF, 1.17 g EDC (6.08 mmol; 1.75 eq.) was added after 60 min. Following the general procedure I 1.64 g tert-butyl (1-oxo-1-(4-(3-(trifluoromethyl)-3H-diazirin-3-yl)phenyl)-5,8,11,14-tetraoxa-2-azahexadecan-16-yl)carbamate (2.99 mmol, 86%) was obtained as an colorless oil.
1H NMR (300 MHz, DMSO) δ 8.69 (t, J = 5.5 Hz, 1H), 7.99 – 7.92 (m, 2H), 7.42 – 7.33 (m, 2H), 6.75 (t, J = 5.8 Hz, 1H), 3.58 – 3.29 (m, 18H), 3.04 (q, J = 6.0 Hz, 2H), 1.36 (s, 9H). 13C NMR (75 MHz, DMSO) δ 165.2, 136.0, 130.2, 128.1, 126.4, 77.6, 69.8, 69.7, 69.6, 69.5, 69.2, 68.8, 54.9, 28.2.

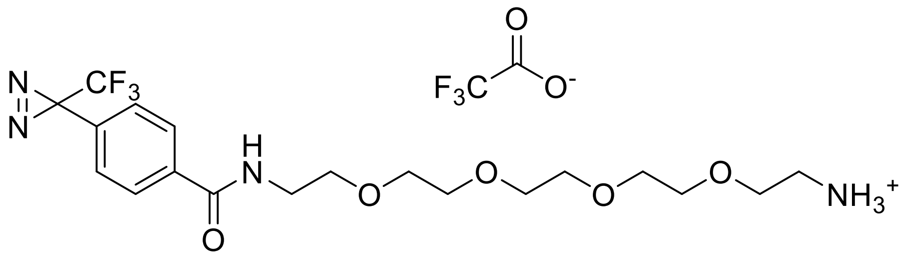


PEG-4 linker **N-(14-amino-3,6,9,12-tetraoxatetradecyl)-4-(3-(trifluoromethyl)-3H-diazirin-3-yl)benzamide:** Starting with 1.62 g tert-butyl (1-oxo-1-(4-(3-(trifluoromethyl)-3H-diazirin-3-yl)phenyl)-5,8,11,14-tetraoxa-2-azahexadecan-16-yl)carbamate (2.95 mmol; 1.00 eq.) in 10 ml dichloromethane and using 7.39 ml TFA (18.9 mmol; 32.5 eq.), following general procedure II N-(14-amino-3,6,9,12-tetraoxatetradecyl)-4-(3-(trifluoromethyl)-3H-diazirin-3-yl)benzamide (1.51 g, 2.69 mmol, 91%) was obtained as TFA salt and colorless oil.

1H NMR (300 MHz, DMSO) δ 8.70 (t, J = 5.6 Hz, 1H), 8.00 – 7.90 (m, 2H), 7.78 (s, 3H), 7.43 – 7.34 (m, 2H), 3.62 – 3.46 (m, 15H), 3.42 (q, J = 5.6 Hz, 2H).

#### 3. Preparation of photoactive cellulose membranes

For activation cellulose membranes were incubated in a solution of sodium hydroxide in water (w(NaOH) = 10%) for 18 hours. Afterwards the membranes were rinsed with ethanol and stored under ethanol until further use for up to three days.

The activated cellulose membranes were taken out of the ethanol and dried at 80 °C in an air stream. The membranes were oxidized in a 2,2,6,6-Tetramethylpiperidinyloxyl (TEMPO)-mediated method with sodium hypochlorite. Therefore, the membranes were incubated in a freshly prepared oxidation solution of 60.9 mg TEMPO (0.39 mmol; 0.2 eq.), 1.45 g NaBr (14.1 mmol; 0.83 eq.) in 292 ml water and 8.48 ml of $NaOCl_{aq}$ (17.0 mmol; c(NaOCl) = 2,00 mol/l; w($Cl_2$) ≈ 12%). The solution was set to a pH-value of 10 with HCl (c = 1 mol/l). The pH-value was checked once again after the membranes were added and adjusted again if necessary. The oxidation was quenched after 60 minutes by rinsing the membranes with ethanol, continued with washing with water.

For NHS-activation a solution of 3.45 g NHS (n = 30 mmol; c = 0.10 mmol/l and 23.0 g EDCl (c = 0.4 mol/l) in 300 ml sodium acetate buffer (pH 5) was prepared. The oxidized membranes were incubated for one and a half hours in the NHS-solution. Afterwards the membranes were taken out and each one was washed with 400 ml of water. The NHS-activated cellulose (NAC) membranes were dried between two paper filters for 18 hours.

Triplicates of NAC membranes (Scheme 1a, **3**) and triplicates of control membranes (Scheme 1a, **1**, **2**) were incubated in a solution of PEG-2 linker (Scheme 1a, **3'**, 10 mM, THF) for 21 hours at room temperature in the dark and under slight shaking of the solution. The membranes were then rinsed in THF (HPLC grade), washed in water for 15 minutes and blocked in a solution of ethanolamine (EA, 1 M, aq.) in the dark. Then the TPD-functionalized cellulose (TFC) membranes were rinsed in water and dried for one hour (30 °C, dark).

#### 4. Photoimmobilization of small molecules

The dried TFC and control membranes were then spotted ten times with 0.5 µl of compound solutions (10 mM, DMSO HPLC grade) for each sample spot using gel loader tips. After evaporation of the solvent *in-vacuo* overnight (RT, dark) each side of the membranes was irradiated with UV-light (365 nm, back and front side) for 30 minutes each. Membranes were rinsed in ethanol, washed intensively in organic solvents (EtOH, DMF, THF, EtOH, water) for one hour each and dried *in-vacuo* overnight at room temperature.

#### 5. Target Pull-Down from whole cell lysates

Jurkat cells were grown in label-free cell culture medium (RPMI-1640, 10% Fetal Bovine Serum (FBS)) and lysed with ice-cold lysis buffer (50 mM HEPES at pH 7.6, 150 mM NaCl, 1 mM EGTA, 1 mM $MgCl_2$, 0.5% Nonident P-40, 0.05% SDS, 0.25% Sodium-deoxycholate) subjected with protease inhibitor (Complete mini tables, Roche) and benzonase. Protein

concentration of the lysate was measured using a detergent compatible protein assay (DC protein Assay, Bio-Rad) and adjusted to 4mg/ml.
Dried cellulose membranes containing the arrayed small molecules were then conditioned for 15 minutes in a LC-MS-grade wash buffer (50 mM HEPES, 150 mM NaCl, 1 mM EGTA at pH 8, 1 mM $MgCl_2$) at pH 7.6 and room temperature. Each membrane was then incubated with Jurkat cell lysate at 4 °C for two hours followed by washing with LC-MS-grade wash buffer three times for 5 minutes also at 4 °C. Membranes were air dried, cellulose spots corresponding to individual photocrosslinked compounds were excised using a paper puncher (d = 4 mm) and transferred into 96-well plates containing LC-MS grade denaturation buffer (6 M urea, 2 M thiourea, 10 mM HEPES, pH 8). Each sample was then treated with dithiothreitol (10 mM, 50 mM ammoniumbicarbonate (ABC), LC-MS grade) for 30 minutes and afterwards with iodacetamide (55 mM, 50 mM ABC, LC-MS grade) for 20 minutes in the dark. Samples were then predigested with LysC (0.5 µg/µl, LC-MS grade) for one and a half hours, diluted four times with ABC (50 mM, LC-MS grade) and digested with trypsin (0.5 µg/µl, LC-MS grade) overnight. Digestion was stopped by reducing the pH (pH < 2.5) using a trifluoroacetic acid solution (10%, LC-MS grade water). Samples were then desalted using StageTip purification (C18, reverse phase, Empore) and stored at 4 °C until further use.

#### 6. LC-MS data acquisition and data analysis

Samples were eluted from stage tips using an elution buffer (0.1% formic acid, 50% acetonitrile in water, LC-MS grade), the solvent was evaporated and samples were resuspended in LC-buffer A (3% acetonitrile, 0.1% formic acid (FA) in water (LC-MS grade). The resulting peptide solutions of all 21 samples (4 compounds and 3 controls, in triplicates) were then analyzed by high resolution LC-MS/MS on a Q Exactive HFX mass spectrometer connected to a nLC1200 system using a short gradient and data dependent acquisition (45 min, DDA: Top20, MS2-resolution: 15K, column: 1.9 µm). The acquired spectra were analyzed in MaxQuant (MQ version 1.6.3.3) using a protein and peptide FDR of 1%, label-free-quantification (LFQ), match-between-runs, re-quantify and MQ standard parameters. This resulted in 3,383 identified proteins for all 21 samples and lysate data combined. Potential contaminants, only identified by side and reverse hits were filtered out, the LFQ data was log2-transformed and replicates grouped together. The data was then filtered on valid values (minimum 3 in at least one group) and missing values were imputed from normal distribution (width: 0.3, down shift: 1.8).
To identify proteins interacting specifically with a given immobilized compound we performed multiple sample testing on the one hand and on the other hand we evaluated the data with a Student's t-test.
First multiple sample testing was performed using LFQ intensities (ANOVA, permutation-based FDR: 5%, 250 randomizations) and ANOVA significant hits were Z-scored and clustered hierarchically. Second protein abundances in the three replicates for a given compound were compared to all other samples using the Student's t-test. Identified proteins with a t-test p-value < 0.01 and fold changes with at least 4 were identified as specific binders. Data filtering and statistical integration was performed in Perseus version 1.6.7.0. Hierarchical clustering was performed in R version 4.1.1.

#### 7. Sample preparation for XPS, ToF-SIMS and ATR-FTIR data acquisition

To evaluate the functionalization of cellulose membranes XPS spectra were acquired after each functionalization step described in 3 (Scheme 1a, **1**, **2**, **3**, **4**, **5**). For preparation of TFC membranes (Scheme 1a, **5**), NAC membranes (Scheme 1a, **3**) were produced as described earlier and incubated in a solution of the photoactive TPD construct (Scheme 1a, **3'**, 10 mM, THF) for 24 hours in the dark. The membranes were then rinsed in THF, washed in THF (15 min, dark) and in water (15 min, dark). Subsequently the membranes were blocked in an ethanolamine solution (EA, 3 M, pH 9.0) twice for an hour and then rinsed and washed twice in water (5 min). The membranes were dried overnight at room temperature in the dark and shipped in falcon tubes.

#### 8. XPS data acquisition

XPS measurements were performed using a K-Alpha+ XPS spectrometer (ThermoFisher Scientific, East Grinstead, UK). The Thermo Avantage software was used for data acquisition and processing. All samples were analyzed using a microfocused, monochromated Al Kα X-ray source (400 µm spot size). The K-Alpha+ charge compensation system was employed during analysis, using electrons of 8 eV energy, and low-energy argon ions to prevent any localized charge build-up. The spectra were fitted with one or more Voigt profiles (BE uncertainty: ± 0.2 eV) and Scofield sensitivity factors were applied for quantification.[2] All spectra were referenced to the C 1s peak (C-C, C-H) at 285.0 eV binding energy controlled by means of the well-known photoelectron peaks of metallic Cu, Ag, and Au, respectively.

#### 9. ToF-SIMS data acquisition

For the assessment of the photoimmobilization of small-molecules we performed ToF-SIMS measurements using cyclosporine A (CsA) as test compound. To that end NAC membranes (Scheme 1a, **3**) and TFC membranes (Scheme 1a, **4**) were spotted in duplicates 10 times with 0.5 µl of a CsA solution (1 mM, DMSO) and dried overnight. One replicate of each was then irradiated with UV-light at 365 nm from both sides for 30 minutes each, whereas the second replicate was kept in the dark. All membranes were then rinsed in EtOH, washed in different organic solvents (EtOH, DMF, THF, EtOH, water) for one hour each and incubated in MeOH overnight in the dark. The membranes were dried and stored at room temperature until further use. The membranes were shipped in falcon tubes protected from light.
*ToF-SIMS* was performed on a TOF.SIMS5 instrument (ION-TOF GmbH, Münster, Germany) equipped with a Bi cluster primary ion source and a reflectron type time-of-flight analyzer. Some samples were slightly outgassing, hence UHV base pressure during analysis was < $2 \times 10^{-7}$ mbar. For high mass resolution the Bi source was operated in bunched mode providing short $Bi_3^+$ primary ion pulses at 25 keV energy, a lateral resolution of approx. 4 µm, a target current of 0.35 pA at 10 kHz repetition rate and 1.1 ns pulse length. For each sample three spots of 500×500 µm$^2$ were analyzed, scanning 128×128 pixel with 75 scans (100 or 125 µs cycle time). Thereby the primary ion dose density was below the quasi static limit ($2 \times 10^{11}$ ions/cm$^2$). Charge compensation was performed by applying a 21 eV electron flood gun and tuning the reflectron accordingly. Peak broadening due to the roughness of the samples and slightly uneven charging was observed. Mass scale calibration was based on hydrocarbon signals. For pos. polarity spectra $C^+$, $CH^+$, $CH_2^+$, and $C_2H_3^+$ were used, for neg.

polarity $C^-$, $CH^-$, $CH_2^-$, $C_2^-$, and $C_3^-$. Original data files, spectra and meta data are available on Radar4KIT.

#### 10. ATR-FTIR data acquisition

ATR-FTIR spectra were recorded on a Bruker Tensor 27, equipped with a platinum ATR-Unit with diamond crystal. The spectra were recorded on a RT-DLaTGS detector with 32 scans and measured against air as background. To follow the functionalization of the cellulose we performed ATR-measurements for each single step. After oxidation a signal at 1600 $cm^{-1}$ for the formed acid is detected (Supporting Figure 1). After activation with NHS we observed two typical signals at 1705 $cm^{-1}$ and 1539 $cm^{-1}$ for the amide group. We see a decrease in the signal at 1705 $cm^{-1}$ when we attach the linker and NHS was substituted. After blocking the non-reacted NHS with ethanolamine (EA) the signal at 1705 $cm^{-1}$ disappeared.

### Supporting Figures

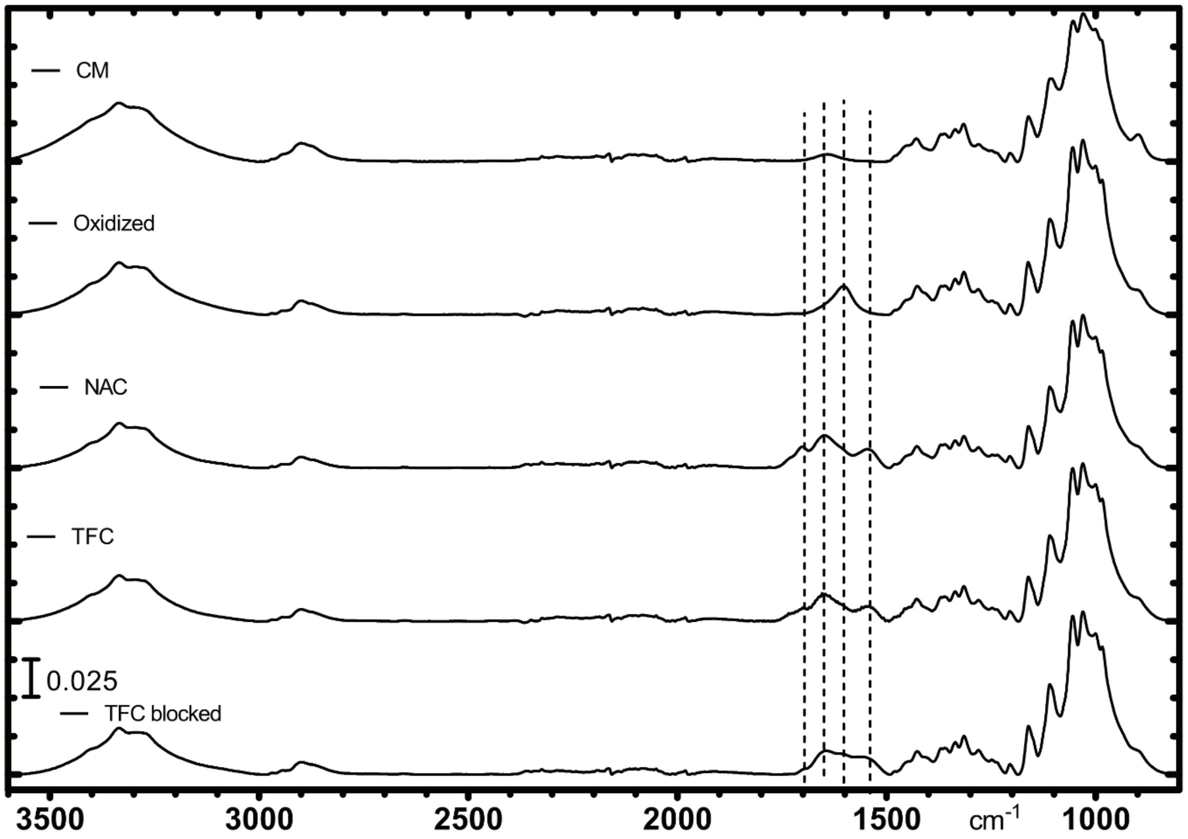


*Supporting Figure 1: ATR-FTIR image of unmodified and functionalized cellulose*

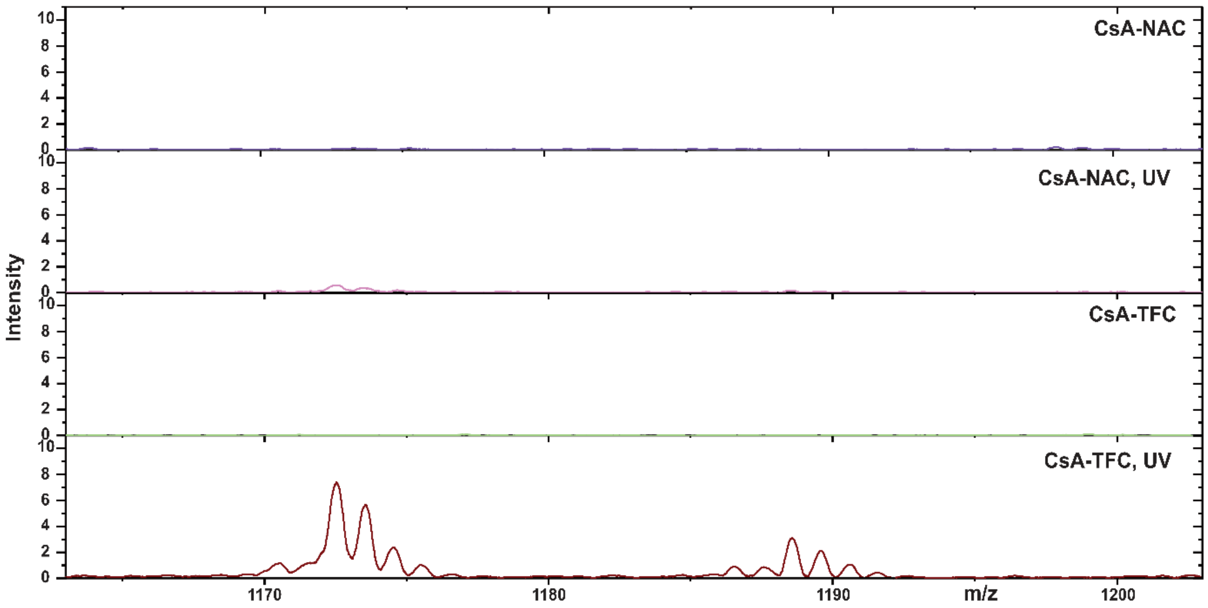


*Supporting Figure 2: smoothed (Savitzky-Golay) ToF-SIMS mass spectra in mass range m/z 1163-1203 of different CsA-spotted cellulose surfaces. TFC: TPD-functionalized cellulose, NAC: NHS-functionalized cellulose, UV: UV-irradiated (365 nm). The $C_{60}H_{106}N_{11}O_{12}$-signal was only significantly observed for the CsA- and UV-irradiated TDB-functionalized cellulose (CsA-TFC, UV).*
